# Using Engineered *Escherichia coli* to Synthesize Squalene with Optimized Manipulation of Squalene Synthase and Mevalonate Pathway

**DOI:** 10.1101/2020.10.28.360032

**Authors:** Chenhao Sun, Yuancheng Ding, Bingjing Cheng, Yeqing Zong

## Abstract

Squalene is the metabolic precursor of sterols and naturally synthesized in the deep-dea shark liver and human sebum. The utilization of squalene is wide such as food, cosmetical, and pharmaceutical industries. This experiment used engineered Escherichia coli to construct the gene circuit for the biosynthesis of squalene. Human squalene synthase (hSQS) efficiently catalyzes the synthesis of squalene. Also, mevalonate (MVA) pathway would increase the yield of the precursor of squalene, farnesyl diphosphate, which then increased the yield of squalene. Meanwhile, the regulation of MVA pathway via different inducer IPTG concentrations and creation of selection pressure by antibiotics were investigated.

## Introduction

Squalene(2,6,10,15,19,23-hexamethyltetracosa-2,6,10,14,18,22-hexaene) is an acyclic triterpenoid, commonly as the metabolic precursor of steroids and hopanoids. The value of squalene presents in various territories. The antioxidant properties are largely used in cosmetic industry as emollient and moisturizer **(Kohno et al., 1995)**; Its ability to reduce serum cholesterol **(Miettinen&Vanhanen, 1994)**, prevent coronary disease, suppress colon tumor **(Rao et al., 1998)**, and boost immmunity **(Kelly, 1999)** enables its use in health functional food. Therefore, the value of squalene reveals the essential need for squalene extraction.

Traditionally, squalene extraction was obtained from plants and animals. Shark liver was the primary source of squalene since the name “squalene” came from the genus “Squalus” of sharks. However, sharks hunting disobeys the notion of animals and environment protection. Also, shark liver contains similar compounds such as steroids, which makes the purification of squalene more complicated. **(Lei, 2015)** An alternative method to avoid ecological and environmental damages would be the extraction from plants **(Jin et al., 2018)**, but it has disadvantages such as low yield and complex purification procedure **(He et al., 2002)**. This experiment recruits microbes fermentation with the benefits of high yield, short growth cycle, low cost, and less limiting environmental factors. The production is investigated in Escherichia coli. As a pervasive host for metabolic engineering, the E. coli strains are easily cultured, have a short doubling time, and can thrive under a variety of growth conditions. Rapid strain development allowed by these techniques can significantly reduce costs for industrial development. **(Pontrelli et al., 2018)** The genetical manipulation of E.coli is clearly studied so it enhances the reliability to engineer new phenotypes.

In triterpenoid metabolic pathway, squalene synthase (SQS, EC 2.5.1.21) plays a vital role and it is the key enzymes catalyzing synthesis of squalene from metabolic precursor farnesyl diphosphate (FPP). SQS catalyzes reductive dimerization of FPP to produce squalene. The first step of catalysis is the condensation of two molecules of FPP. SQS creates head-to-head orientation for the two molecules to form presqualene diphosphate (PSPP). This process requires metal ion Mg2+ to stabilize the phosphate groups of FPP. **(Davidson, 2007)** The PSPP undergoes heterolysis, then 1,2 sigmatropic rearrangements produces a cyclopropyl intermediate. The second step of catalysis requires NADPH to reduce PSPP. The hydride transfer from NADPH causes the cleavage of the cyclopropyl group. The hydrocarbon chain then undergoes rearrangement to form squalene, with side products: PPi, NADP+, and H+. **(Tansey&Shechter, 2000)** The final step is the release of squalene from SQS.

To optimize the yield of squalene, different sources and the identification and removal of domains were studied. SQS variants from bacterial (Trypanosoma) and human sources were compared and engineered in E.coli, which reveals the truncated human squalene synthase (hSQS) investigated shows highest yield. Besides the high yield, compared with Trypanosoma squalene synthase (tSQS) variants, hSQS produced less dehydrosqualene **(Katabami et al., 2014)**, which is an unwanted molecule that complicates purification process. Therefore, hSQS is more acceptable in production of squalene, and the variant ensures the expression in E.coli results in active hSQS enzyme.

SQS catalyzes FPP into squalene, the synthesis of reactants FPP is a notable step to increase the yield of squalene. The biosynthesis of FPP is mainly through two pathways: methylerythritol phosphate (MEP) pathway and mevalonate (MVA) pathway. MEP pathway is mainly found in plants, bacteria and alga (Hale&O’Neill, 2012), while MVA pathway occurs in the cytosol of eukaryotes, archaea and higher plants **(Lee&Schimidt-Dannert, 2002)**. Therefore, MEP pathway is endogenous but unengineered E.coli do not contain MVA pathway metabolism. With engineered MVA pathway, the two pathways would both work in the E.coli, which enhances the yield of precursor FPP (Figure 1).

**Figure 1.**
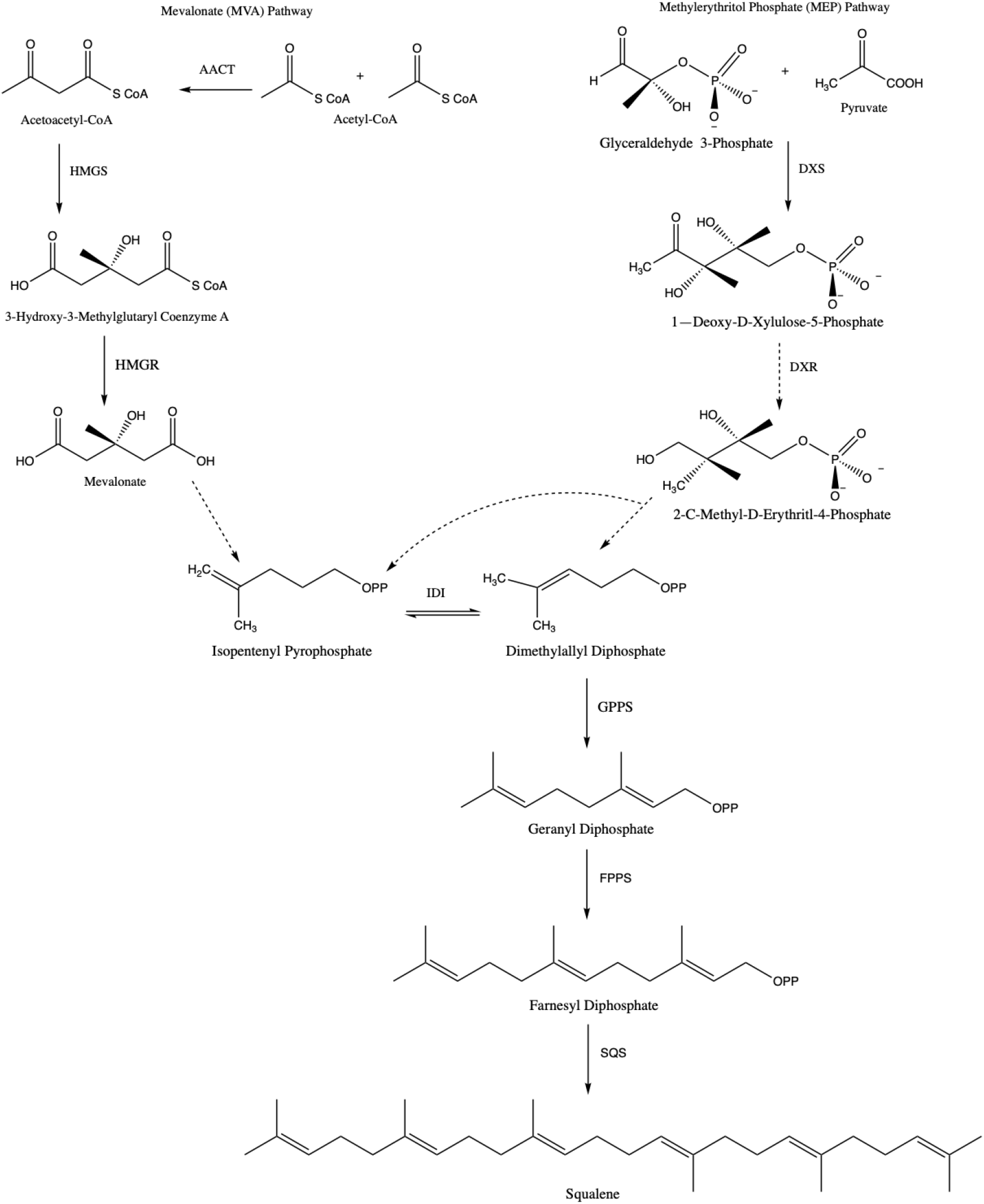
Endogenous and Synthetic pathways for squalene synthesis. AATC: Acetoacetyl-CoA thiolase; HGMS: Hydroxymethylglutaryl-CoA synthase; HMGR: Hydroxymethylglutaryl-CoA reductase; DXS: 1-deoxy-Dxylulose-5-phosphate synthase; IDI: Isopentenyl diphosphate isomerase; GPPS: Geranyl diphosphate synthase; FPPS: Farnesyl diphosphatesynthase; SQS: Squalene synthase

Furthermore, utilizing inducible expression system is important if squalene is under industrial production. Inducible expression systems are essential molecular tools for production of recombinant proteins in cells, for synthesis and degradation of small molecules catalyzed by the enzymes expressed from the expression system. The inducible lac promoter is one of the most commonly used promoters for heterologous protein expression in *E. coli*. Isopropyl-β-D-thiogalactoside (IPTG) is currently the most efficient molecular inducer for regulating this promoter’s transcriptional activity. **(Briand et al., 2016)** Therefore, the IPTG-inducible expression system was used in the squalene production, and the optimum concentration was investigated.

Also, the use of antibiotics is another aspect to investigate the squalene biosynthesis optimized conditions. Since engineered plasmids are often lost in culture **(Summers, 1998)**, it is essential to impose a selective pressure to ensure plasmid stability, which is often achieved by adding antibiotics to the culture medium. However, industrial production via engineered bacteria does not use antibiotics during fermentation because it is expensive and increases the growth burden for bacteria. Therefore, it is an important factor that would affect the yield of squalene but the economic aspect worth investigation.

## Materials and Methods

### Strains and plasmids

Plasmids pMevT and pMBIS were used from **(Martin et al., 2003)** that enhance FPP synthesis via MVA pathway. Two plasmids were both engineered in Escherichia coli. The human squalene synthase gene (hSQS) refers to the variant in **(Katabami et al., 2014)** with the highest efficiency of production, but the gene was condon optimized in this study. The tested strain was E. coli DH10B. All the maps of plasmid mentioned in this paper were included in the **supplementary material**.

### Reagents

FastPfu High-Fidelity Master Mix was purchased from New England BioLabs (NEB). TIAN Prep Rapid Mini Plasmid Kit was purchased from TIANGEN. The agarose gel extraction purification kit was purchased from TIANGEN. Other reagents and drugs without description are all domestically analytical grade. PCR primer synthesizing and sequencing were provided by Shenzhen Qingke.

**Table 1.** Reagents and their companies.

| Reagent | Company |
| --- | --- |
| FastPfu High-Fidelity Master Mix | New England BioLabs (NEB) |
| TIAN Prep Rapid Mini Plasmid Kit | TIANGEN |
| The agarose gel extraction purification kit | TIANGEN |

### Medium

Media composition recipes are refered from **(Green&Sambrook, 2012)**

**Table 2.** Preparation of Liquid Luria-Bertani (LB) Medium.

| Chemical Composition | Mass (g) / Volume (L) |
| --- | --- |
| Tryptone | 10 g |
| NaCl | 10 g |
| Yeast extract | 5 g |
| Water | 1 L |

**Table 3.** Preparation of Terrific Broth (TB) Medium.

| Chemical composition | Mass (g) /Volume (L) |
| --- | --- |
| Tryptone | 11.8g |
| Yeast extract | 23.6g |
| KH<sub>2</sub>PO<sub>4</sub> | 2.2g |
| K<sub>2</sub>HPO<sub>4</sub> | 9.4g |
| 80% Glycerol | 0.005 L |
| Water | 1 L |

2.2g of KH2PO4 and 9.4g of K2HP4 were firstly dissolved in 100 mL water, sterilized through 0.2um membrane filtration. 11.8g of tryptone and 23.6g of yeast extract were dissolved in 900mL of sterile water, then 5mL of 80% glycerol was added. Then this solution was sterilized for 25 minutes in an autoclave at 115°C. When it was cooled to less than 60°C, phosphate solution was added to this solution.

### Transformation

50μL DH10B competent E.coli strain were taken and thawed slowly on ice. The DNA (plasmids or assembled products) that would be transformed was added and flicked to mix with competent cells. It was then placed in ice for 30 minutes. The reaction products were put in the 42°C water bath for 60s heat shock and taken ice-bath immediately for 3 minutes. During this process, EP tubes were not moved. 300μL antibiotic-free LB liquid medium was added and cultured in a 37°C shaker at 220rpm for 60 minutes.

For plasmid transformation, 100μL of resuspended bacteria solution was absorbed and coated on a solid LB plate containing the corresponding antibiotic, which was inverted and cultured at 37°C for 16 hours; For the transformation of lassembled products, resuspended bacterial solution was centrifuged at 4000 RPM for 1 minute. 250μL supernatant was removed. The remaining liquid medium was blown to mix with the precipitation and coated on a solid LB plate containing the corresponding antibiotic, which was then inverted and cultured at 37°C for 16 hours.

### Colony PCR

The primers, ddH2O and Mix were advancely thawed. The PCR reaction system was prepared according to **Table 4** and blown with a pipette. The reaction system was then thoroughly mixed on a vortex oscillator and placed on the ice.

**Table 4.**
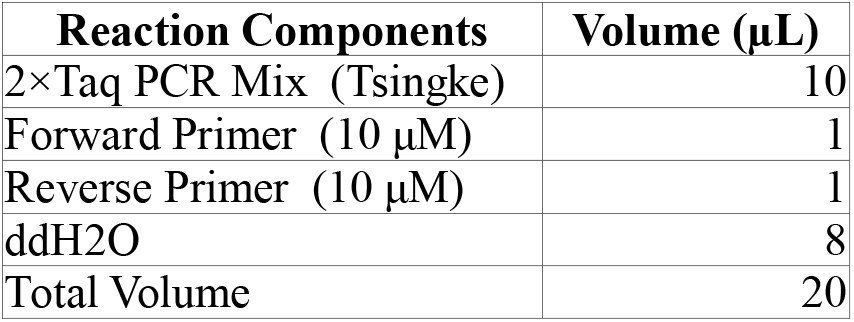
PCR Preparation System.

The corresponding resistant solid LB plate was prepared. The PCR tube and plate were marked with marker pen for one-to-one correspondence. The ultraviolet lamp of the clean bench was advancely turned on to release ozone for sterilization. Then the lighting and air supply near the burning alcohol lamp were turned on. The single colony was picked by the sterilized toothpick. A line was drawn by this toothpick on the LB plate as the copy of the original colony. The toothpick was then inserted and gently rotated in the PCR tube with corresponding numbers and thrown into the trash can. The last step was repeated to obtain multiple potential single colonies. The PCR tube was put into the PCR thermal cycler with the procedure program shown in **Table 5**.

**Table 5.** PCR Reaction Procedure.

| Step name | Temperature(°C) | Duration | Cycle number (cycle) |
| --- | --- | --- | --- |
| Predenaturation | 94 | 10min | 1 |
| Denaturation | 94 | 30s | 30 |
| Annealing | 55 | 30s | 30 |
| Degeneration | 72 | 1kb/min | 30 |
| Extension | 72 | 5min | 1 |

PCR products were taken out and 1% agarose gel was used for DNA length detection. If bands were single and bright, the products would be used for purification.The plasmid with the correct length could be used in the following experiments.

### Fermentation

#### Effects of Different IPTG Concentration on Bacterial Growth

Under the same culture conditions, different concentrations of IPTG inducers were added to conduct fermentation experiments. The phase of bacterial growth was observed and OD value was measured for comparative analysis. Three single colonies were taken from solid LB plate and cultured separately in 3 mL of TB medium for 3 hours at 37°C and 1000 rpm. 0-500 μL of IPTG was added to induce and culture for 48 hours.

#### Effect of the Addition of Antibiotics on Squalene Production

Group A: Antibiotics (final concentration: tetracycline 5 g / mL, chloramphenicol 25 g / mL, kanamycin sulfate 50 g / mL) were added.

Group B: No antibiotics.

#### Extraction of Squalene

After fermentation, the fermentation liquid was centrifuged at 10000 rpm for 1 minute. The precipitation was remained and then blown with 1 mL pf 1% (W/V) normal saline to resuspend. Centrifugation was repeated at the same speed for 1 minute. The supernatant was discarded, while the precipitation was washed again. Then 100 μL normal saline was added and the precipitation was resuspended. 1 ml of acetone was added to lyse the cells. The contents were secreted, which were shaken gently and allowed to stand at room temperature for 2 minutes. After centrifugation at 10000rpm for 1 minute, the supernatant was transferred to a clean EP tube for further detection.

#### Analysis of UPLC

Instrument: Shimazu LC-20A.

Stationary phase: GIST-HP (3 μm C18, 2.1×50 mm).

Mobile phase: 100% methanol.

Flow rate: 0.3ml / min.

The sample of squalene was diluted with acetone to obtain the standard for liquid phase detection with the sample. The curve of concentration peak area of the standard solution of squalene was drawn. Whether there were products generated was generated according to the peak time and the yield of product was calculated according to the peak area (Figure 2).

**Figure 2.**
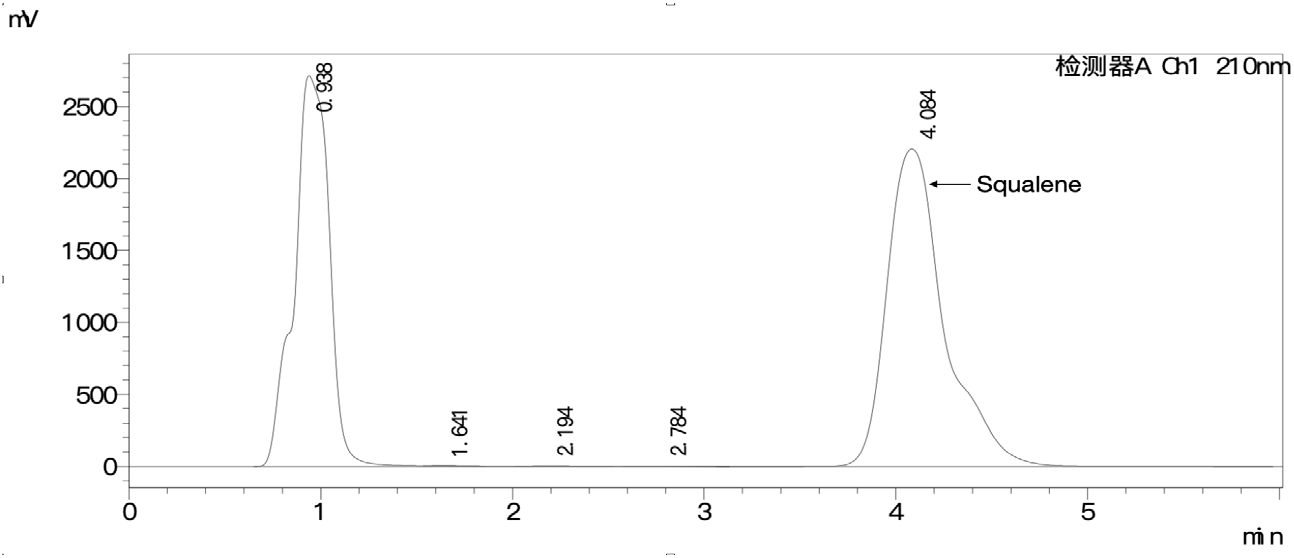
UPLC Traces of Squalene.

## Results

### Plasmids Design and Construction

Three plasmids that were pTYT-hSQS, pMBIS, and pMevT were constructed and transformed into E.coli DH10B for squalene production.

#### pTYT-hSQS

The pTAC-RiboJ-lacZα-rrnBT1 fragment was obtained and amplified from pRG **(Zong et al., 2017)** via PCR. A high-copy IPTG-inducible vector pTYT was constructed via Gibson Assembly of the PCR-amplified fragment and pET-28b vector. Then the hSQS gene sequence was inserted into pTYT via Golden Gate Assembly (Figure 3).

**Figure 3.**
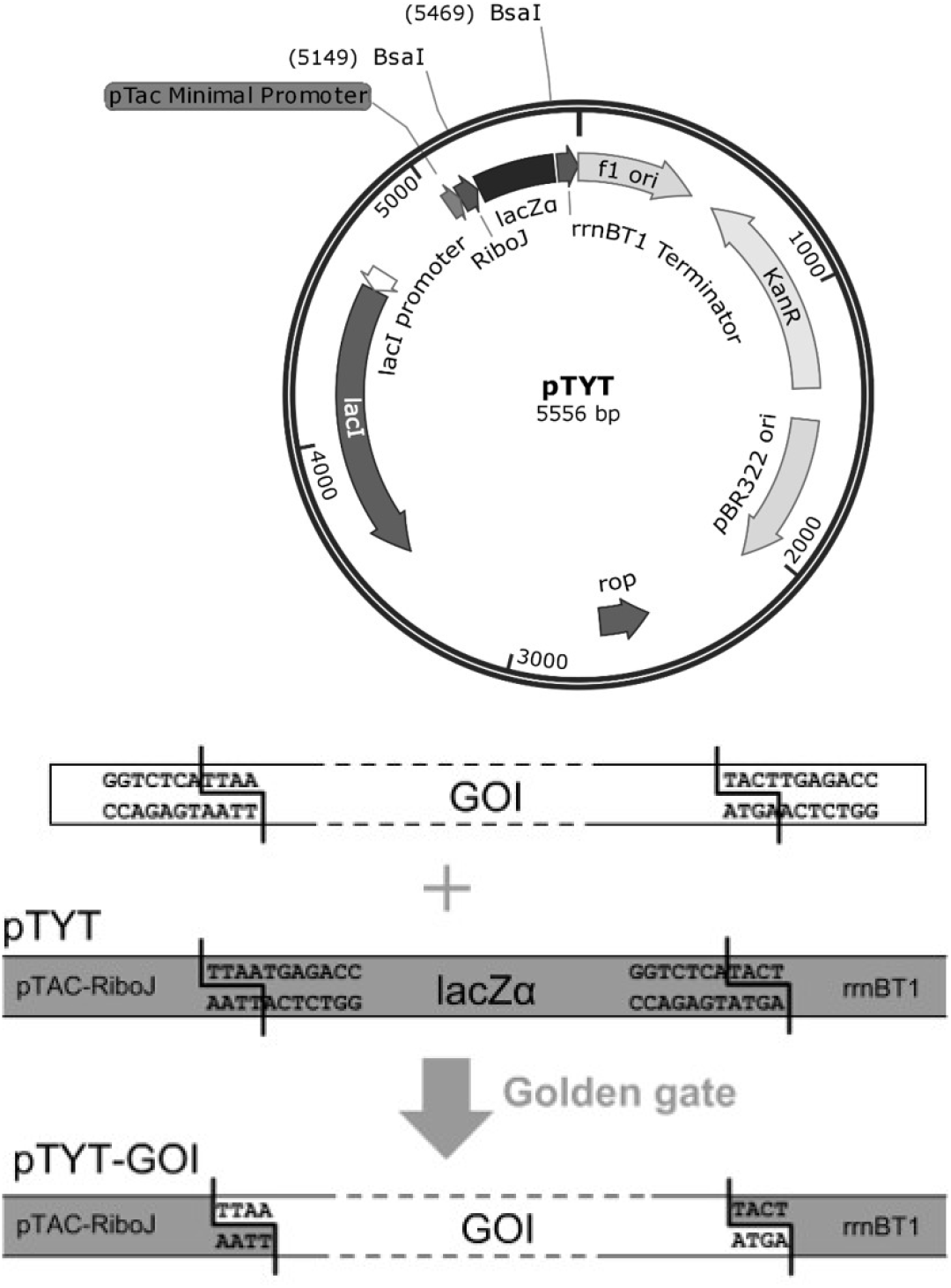
Construction of squalene expression system.

#### pMBIS and pMevT

These two plasmids with genes encoding for enzymes involved in MVA pathway were referred from **(Martin et al., 2003)**.

Firstly, the vector plasmid pTYT was constructed to express SQS gene. The pTYT plasmid is provided by two open plasmids. The part containing the resistance gene, replication initiation site and lacI expression unit was derived from the pET-28b (+) plasmid, while the part containing the pTAC promoter, RiboJ spacer, BsaI restriction site, lacZ α marker and Terminator was derived from the pRG plasmid. LacZ α fragments could be replaced by one or more gene fragments by Goldengate assembly method. The inserted gene fragment was controlled by pTAC strong promoter and could be induced by IPTG. In this project, the hSQS genome containing RBS on the pTYT plasmid was loaded.

### Cells Growth under Different Conditions

As the MEP pathway of E. coli could produce the substrate FPP of hSQS, the squalene could be detected when pTYT-hSQS was transformed directly into the host cell of Escherichia coli. Due to the strong leakage expression of pTAC promoter, 4.1 ±0.1mg/L squalene could be detected in the culture without induction in this experiment. The production of squalene should be increased by increasing the substrate supply of hSQS, so pMevT and pMBIS plasmids from Keasling laboratory were transferred into the strain, which affected the heterologous MVA pathway to be constructed in the strain to enhance the supply of FPP. Under the same culture conditions, the yield of squalene of the strains with MVA pathway was more than double that of the strains without MVA pathway. On pMevT and pMBIS plasmids, the enzyme genes involved in the MVA pathway were controlled by the pLac promoter, so it could be imagined that the expression of each enzyme in the pathway could be increased by adding inducer IPTG, which helped increase the final yield. However, when the concentration of IPTG in the medium reached 500 μM, the yield of squalene decreased by nearly half compared with that without induction. The cell density in the culture also decreased significantly at this time. It is worth-noting that when there were only pMevT and pMBIS plasmids in the cell, the addition of inducer had no significant effect on the growth, so this growth inhibition might be the result of the over-expression of hSQS (Figure 4).

**Figure 4.**
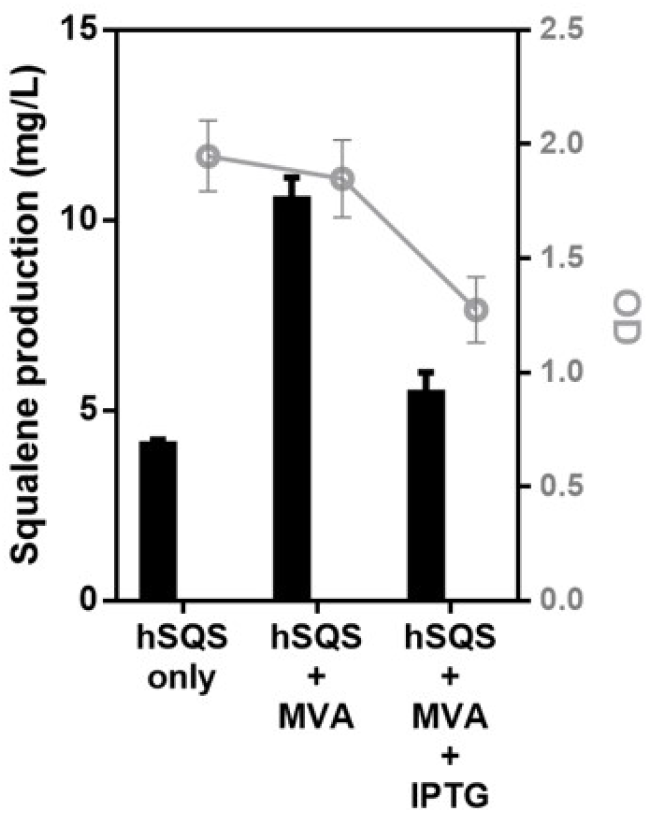
Effects of different treatment conditions on cell growth and squalene production.

#### Squalene Production under Different Conditions

By adjusting the concentration of IPTG, a balance between synthase expression and cell growth can be found, so as to increase the final production of squalene. However, in the gradient induction culture experiment, we found that the cell density decreased significantly when the IPTG concentration was only 20 μ M. although the squalene yield per unit biomass was twice that of the uninduced, the total yield did not increase significantly. At higher concentration of inducer, the yield of squalene decreased further (Figure 5).

**Figure 5.**
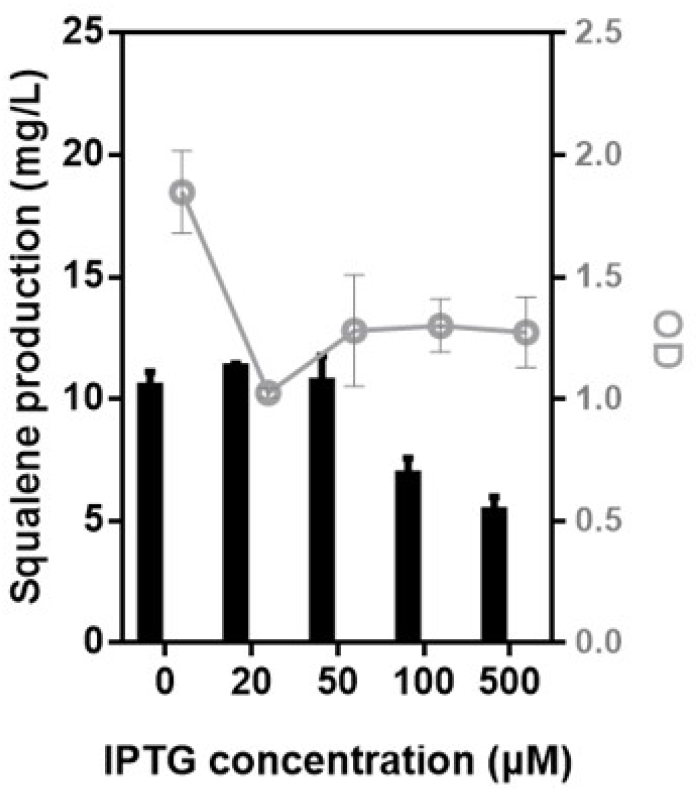
Effects of different IPTG concentrations on cell growth and squalene production after adding antibiotics.

We believe that properly reducing the cell growth resistance may help to increase the cell concentration and then increase the total yield of the product. So in the next experiment, we no longer add antibiotics to the culture medium. The results showed that the cell growth was restored under the condition of no screening pressure, but we noticed that the yield of squalene decreased when the concentration of inducer was less than 50 μ M. We think that the decrease of screening pressure may lead to the loss of plasmids in some strains, thus reducing the yield. But unexpectedly, when the concentration of IPTG was above 100 μ M, the total yield and unit yield of squalene increased compared with the experimental group with antibiotics, reaching 16.0 ±2.0mg/L when the concentration of IPTG was 100 μ M. however, when the concentration of IPTG was above 100 μ M, the total yield and unit yield of squalene were increased compared with the experimental group with antibiotics (Figure 6).

**Figure 6.**
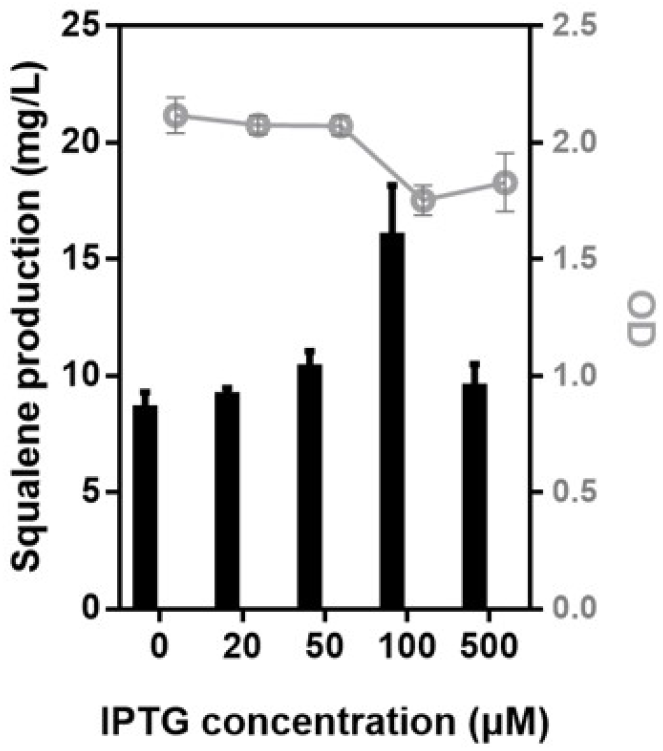
Effects of different IPTG concentrations on cell growth and squalene production in the absence of antibiotics.

## Discussion

In this study, the original Pt7-lac transcription control system based on pET-28b (+) was replaced by the PTC transcription control system. The BsaI enzyme cut-off site contained lacZα markers which allowed it to act as a granule of the Golden Gate assembly response and be assembled in single step with more than one inserted fragment (Engler, Gruetzner et al. 2009).

In previous studies on the production of squalene by introducing hSQS gene into E. coli (Furubayashi, Li et al. 2014, Katabami, Li et al. 2015), the nucleic acid sequence of hSQS was directly derived from human cDNA and was the truncated version at the N-terminal (Thompson, Danley et al. 1998), while this study used the E. coli codon optimization of gene sequence. The experimental results showed that it also had the ability to catalyze and generate squalene in prokaryotic cells, but the yield was low, which might be the result of low supply of FPP. If optimized MVA approach could be adopted, higher yield would be obtained.

For the optimization of culture conditions, it was mainly about seeking for the optimal, quantitative combination of factors such as medium composition, culture temperature, amount and time of inducer addition. Screening pressures such as antibiotics were generally considered to be conducive to maintain the genetic stability of plasmid that contained strains and increase the yield of product. In this study, in addition to the amount of inducers, the addition of antibiotics was also added as a variable for the optimization of culture conditions. Under the condition of high concentration of inducer, the biomass and squalene yield of the experimental group without antibiotics were significantly increased, compared with the experimental group with antibiotics. The mechanism of increasing yield was still unclear. However, it should be noted that pTYT expression vector had an immutable pBR322 replication starting site, while it had a mutable copy number affected by antibiotics or growth pressure, which further affected the expression level of its inserted genes. It might be a clue to explain the experimental phenomenon in this study.

## Supporting information

Plasmids' Maps

